## Additional file 1 for "Extensive genomic and transcriptomic variation defines the chromosome-scale assembly of *Haemonchus contortus*, a model gastrointestinal worm"

### Supplementary information - Figures

**Section 1. Chromosome structure of Haemonchus contortus** **2**
Figure S1. Genome assembly pipeline and improvement timeline. 2
Figure S2. Assembly of the X chromosome. 3
Figure S3. Conservation of ortholog synteny between H. contortus and C. elegans. 5
Figure S4. Visualisation of genome completeness metrics using BUSCO and CEGMA. 7

**Section 2. Resolving haplotypic diversity and repeat distribution within the chromosomes** **8**
Figure S5. Distribution of repetitive units throughout the genome identified with RepeatMasker. 8

**Section 3: Generation of a high-quality transcriptome annotation incorporating short and long reads** **9**
Figure S6. Annotation pipeline schematic used to incorporate RNAseq and IsoSeq data into a single annotation. 9

**Section 4: Transcriptional dynamics throughout development and between sexes** **10**
Figure S7. Transcript co-expression profiles across life stages of Haemonchus contortus. 10

**Section 5: Transcriptional complexity is defined by extensive cis- and trans-splicing** **11**
Figure S8. Comparison of splice leader and splice site sequence diversity 11

**Section 5: Distribution of global genetic diversity throughout the chromosomes** **13**
Figure S9. Principal component analysis (PCA) of mitochondrial diversity 16
Figure S10. Distinct differences between mtDNA diversity in South Africa 17
Figure S11. X chromosome coverage variation 18

**References** **19**

#### Section 1. Chromosome structure of *Haemonchus contortus*

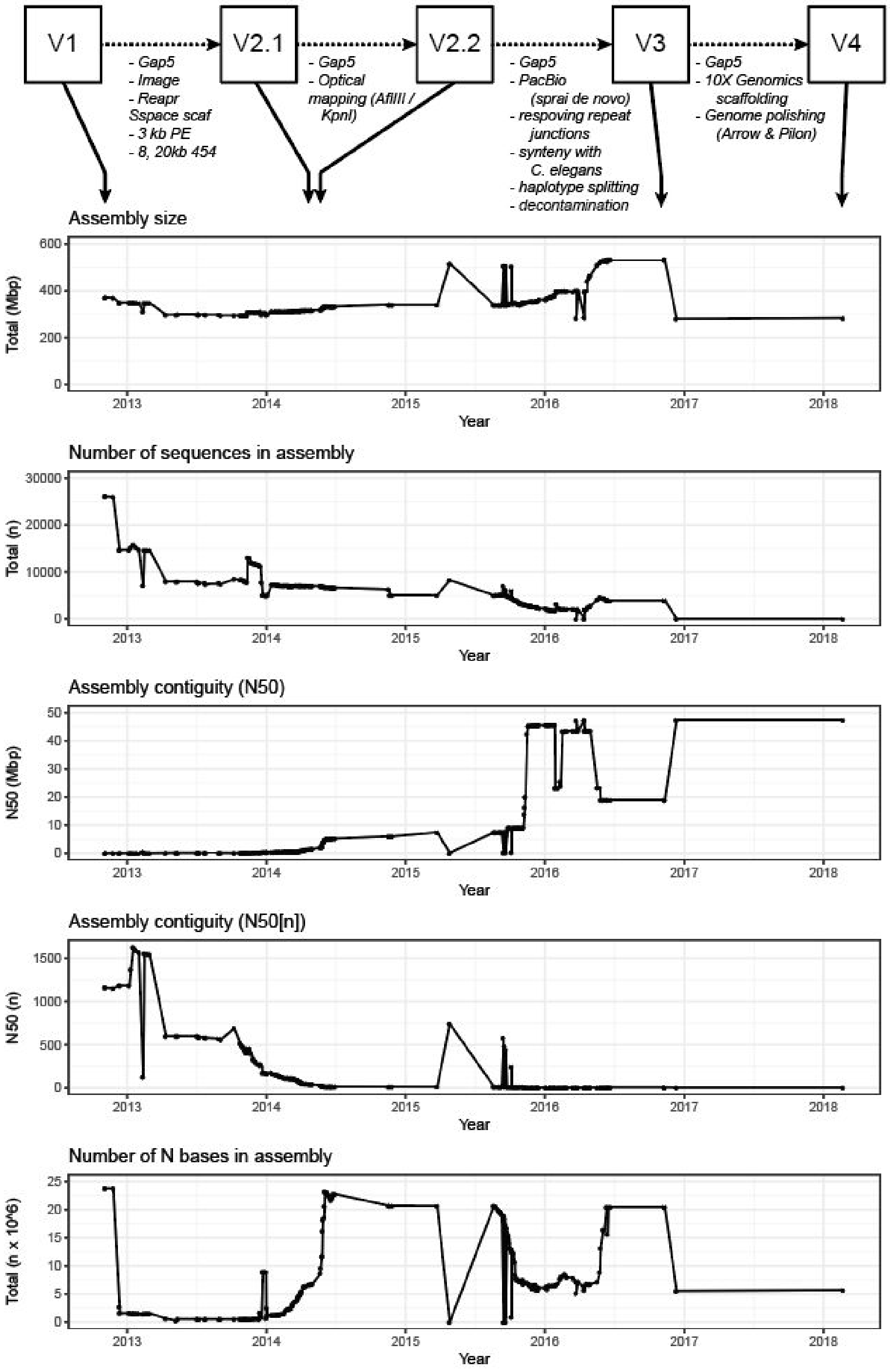


**Figure S1. Genome assembly pipeline and improvement timeline.**
The genome was iteratively improved from the published version in 2013 to the complete genome assembly presented here as new technologies became available. Here, we present the changes in the assembly overtime, including the assembly size, number of contig and/or scaffold sequences, the assembly contiguity measured using N50 and N50(n), as well as the number of N bases representing missing data. As a reference point, the V3 genome was used in the genetic map [1] and ivermectin backcross [2] analyses.

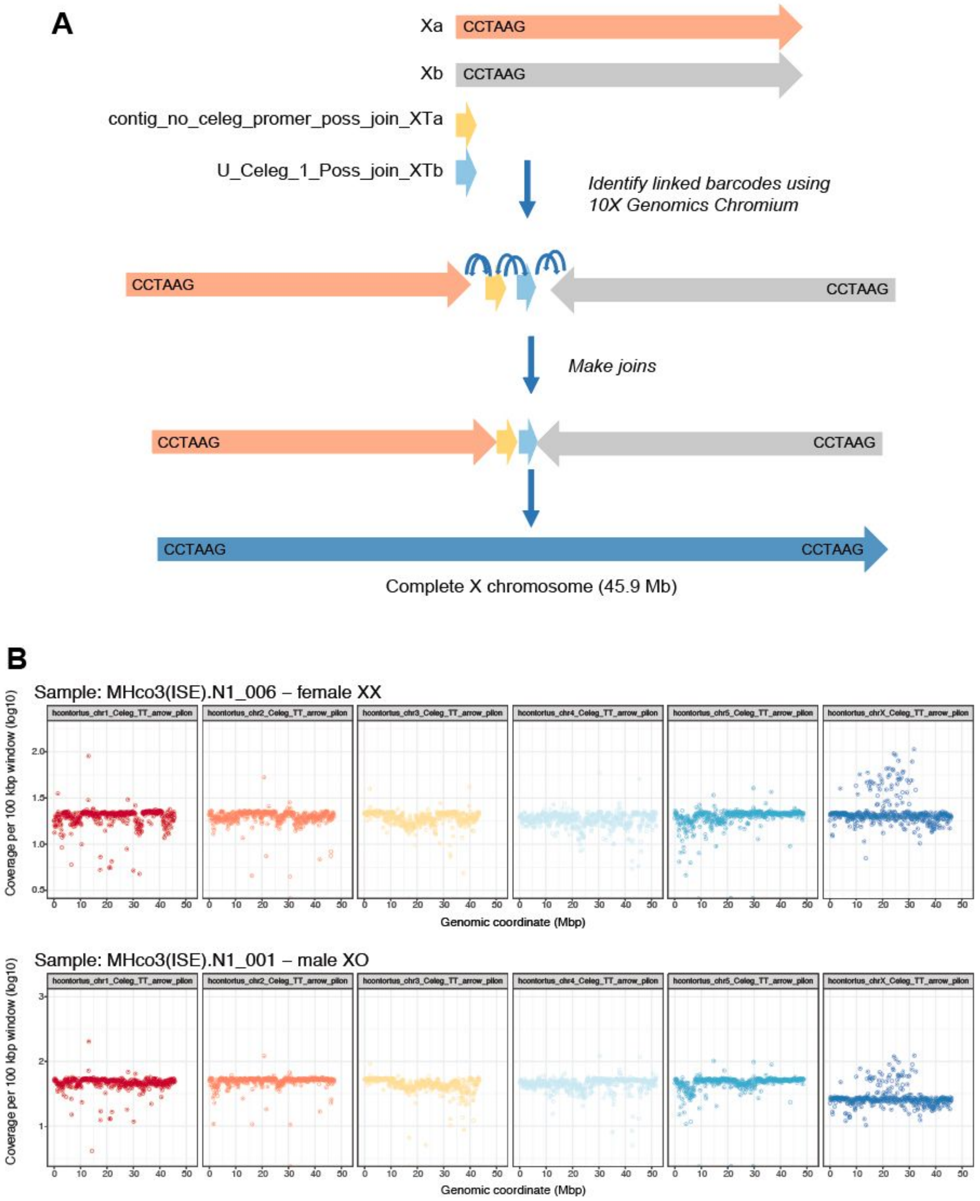


**Figure S2. Assembly of the X chromosome.**

**A.** A major update between the V3 and V4 versions of the genome was the scaffolding of the X chromosome into a single sequence. We used 10X Chromium linked reads together with the ARCS/LINKS scaffolding pipeline [3,4] to join two major scaffolds containing telomeric sequences, and two short, unplaced X-linked scaffolds (putatively linked to X by shared repeats and coverage differences between male and female single worm sequencing), resulting in the single 45.9 Mb sequence. **B.** Genome-wide sequence coverage between a single male and female parasite, illustrating the relative coverage difference of the hemizygous male X chromosome.

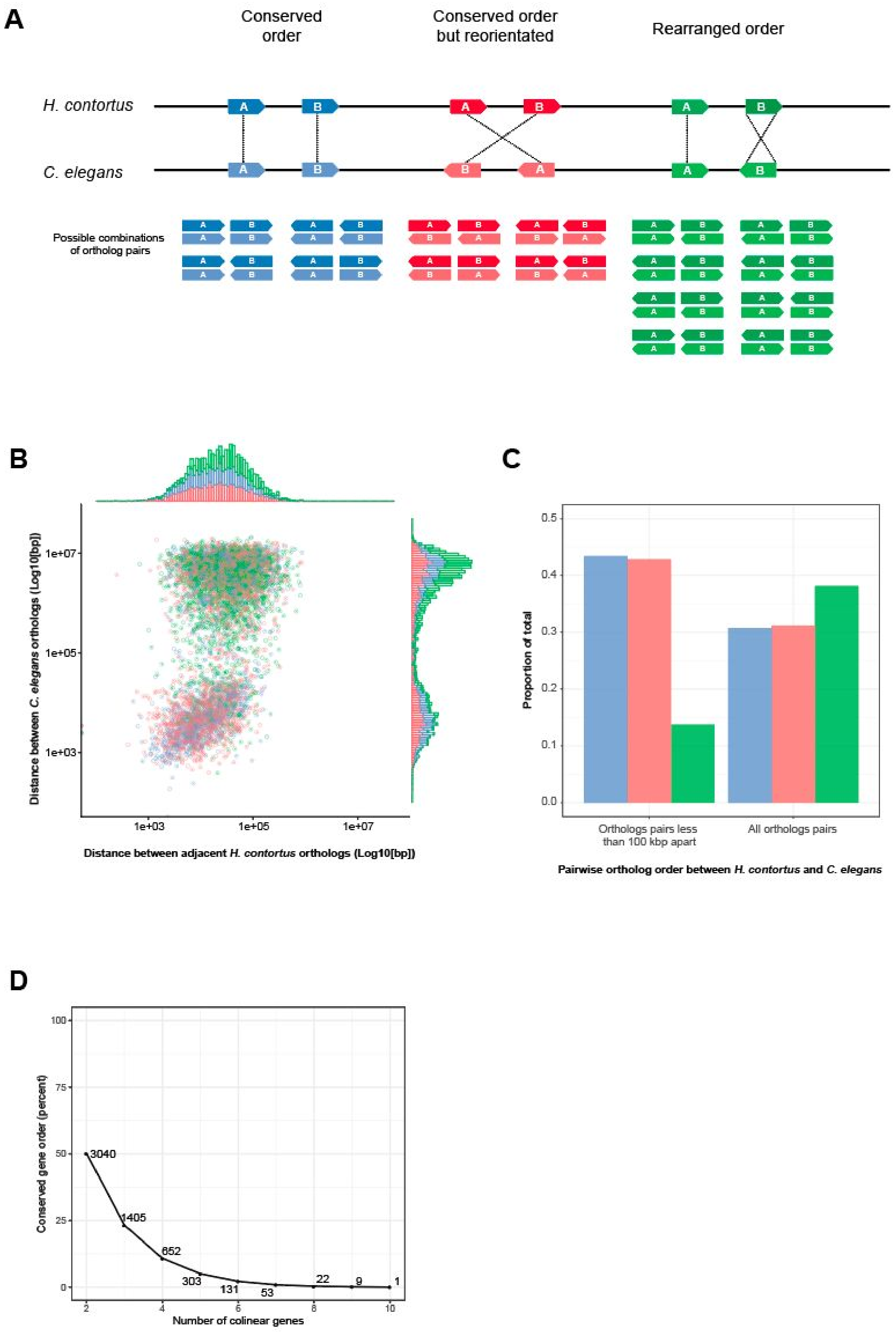
A
Conserved order
Conserved order but reorientated
Rearranged order
H. contortus
C. elegans
Possible combinations of ortholog pairs
B
Distance between C. elegans orthologs (Log10[bp])
Distance between adjacent H. contortus orthologs (Log10[bp])
1e+03
1e+05
1e+07
C
Proportion of total
Orthologs pairs less than 100 kbp apart
All orthologs pairs
Pairwise ortholog order between H. contortus and C. elegans
D
Conserved gene order (percent)
Number of colinear genes
3040
1405
652
303
131
53
22
9
1

**Figure S3. Conservation of ortholog synteny between *H. contortus* and *C. elegans*.**
**A.** Schematic of the arrangement of ortholog pairs analysed. Each comparison throughout compares adjacent genes in *H. contortus* for which a one-to-one ortholog was found in *C. elegans*. The experiment, therefore, examines the proportion of *H. contortus* gene pairs that have the same (or different) orientation in *C. elegans*. "Conserved order" (blue) represents pairs of orthologs that are in the same relative position and orientation between the two species. "Conserved order but reorientated" (red) represents the pairs that are in the same order, but are in a different orientation between the two species, for example, there has been a large-scale structural inversion in one species (but not the other) that changes the orientation of both genes in that species due to the inversion. "Rearranged" (green) represents instances where a change in the orientation of one of the two genes in the pair has occurred in one of the species. **B.** Scatterplot with density histograms per axis comparing the distance between adjacent *H. contortus* gene pairs (x-axis), and the relative distance between the orthologs of those gene pairs in C. elegans (y-axis). Points and bars are coloured as described in **A. C.** Proportion of pairwise groups described in **A**, comparing the frequency of those genes that were found within 100 kb of each other with all gene pairs regardless of their distance. **D.** Comparison of the proportion of colinear orthologs between the two species. Groups of colinear genes, ranging from 2 to 10 genes per group, are shown on the x-axis, and the proportion of genes in that group are shown on the y-axis. The total number of genes within each group are indicated on the plot.

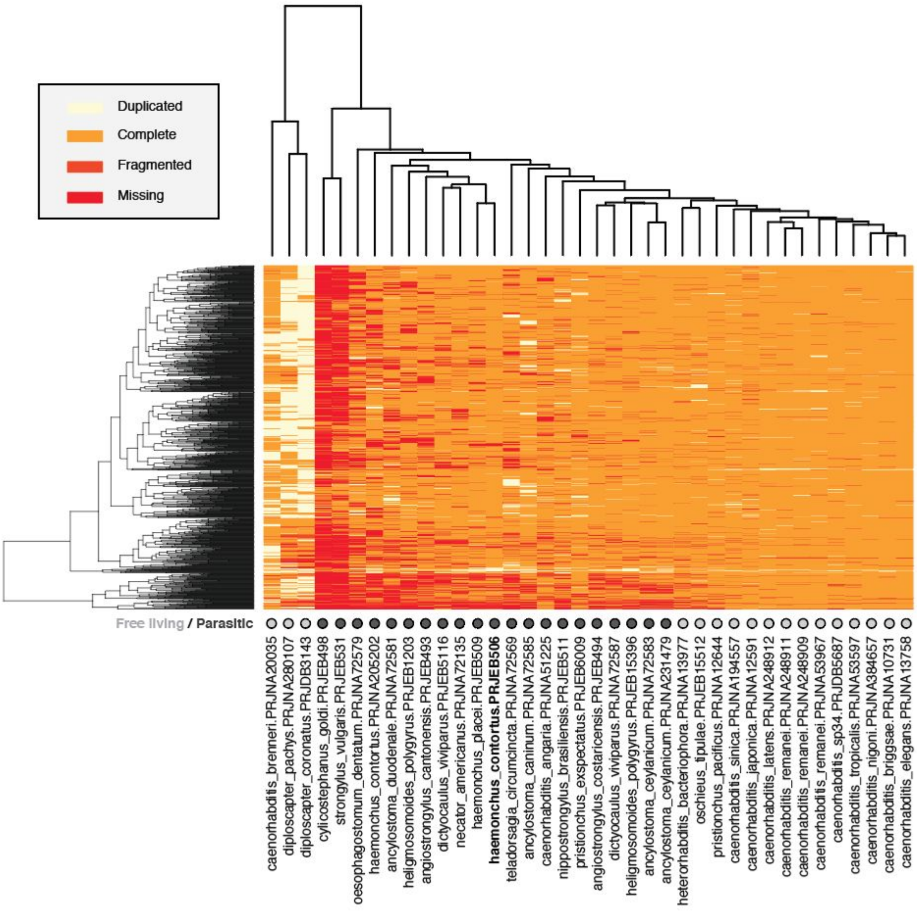
Duplicated
Complete
Fragmented
Missing
Free living / Parasitic
caenorhabditis_brenneri.PRJNA20035
diploscapter_pachys.PRJNA280107
diploscapter_coronatus.PRJDB3143
cylicostephanus_goldi.PRJEB498
strongylus_vulgaris.PRJEB531
oesophagostomum_dentatum.PRJNA72579
haemonchus_contortus.PRJNA205202
ancylostoma_duodenale.PRJNA72581
heligmosomoides_polygyrus.PRJEB1203
angiostrongylus_cantonensis.PRJEB493
dictyocaulus_viviparus.PRJEB5116
necator_americanus.PRJNA72135
haemonchus_placei.PRJEB509
haemonchus_contortus.PRJEB506
teladorsagia_circumcincta.PRJNA72569
ancylostoma_caninum.PRJNA72585
caenorhabditis_angaria.PRJNA51225
nippostrongylus_brasiliensis.PRJEB511
pristionchus_exspectatus.PRJEB6009
angiostrongylus_costaricensis.PRJEB494
dictyocaulus_viviparus.PRJNA72587
heligmosomoides_polygyrus.PRJEB15396
ancylostoma_ceylanicum.PRJNA72583
ancylostoma_ceylanicum.PRJNA231479
heterorhabditis_bacteriophora.PRJNA13977
oschieus_tipulae.PRJEB15512
pristionchus_pacificus.PRJNA12644
caenorhabditis_sinica.PRJNA194557
caenorhabditis_japonica.PRJNA12591
caenorhabditis_latens.PRJNA248912
caenorhabditis_remanei.PRJNA248911
caenorhabditis_remanei.PRJNA248909
caenorhabditis_remanei.PRJNA53967
caenorhabditis_sp34.PRJDB5687
caenorhabditis_tropicalis.PRJNA53597
caenorhabditis_nigoni.PRJNA384657
caenorhabditis_briggsae.PRJNA10731
caenorhabditis_elegans.PRJNA13758

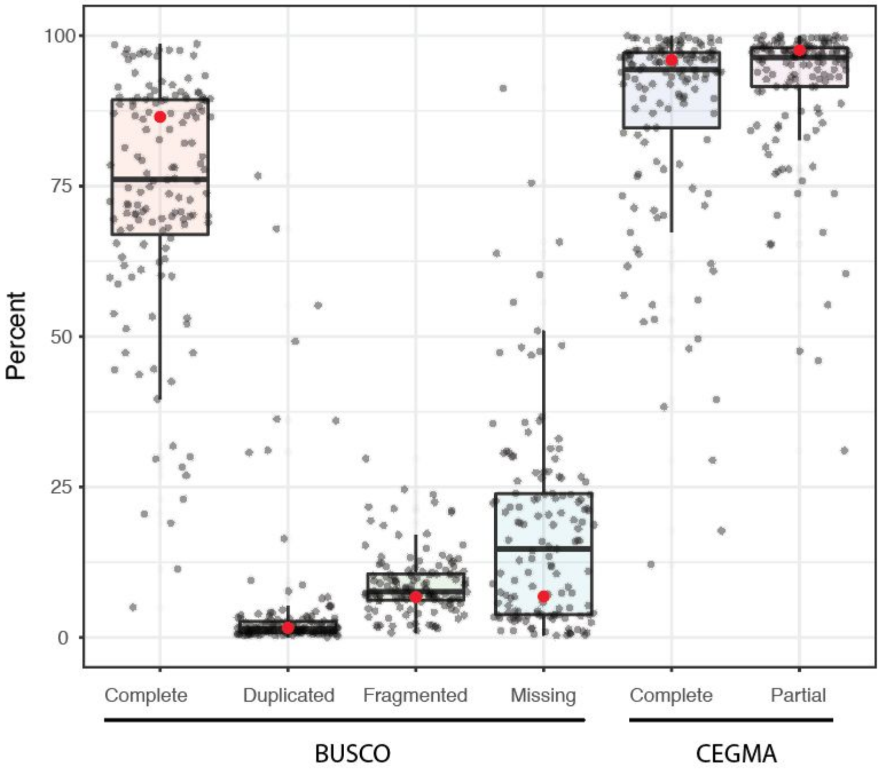
100
75
50
25
0
Percent
Complete
Duplicated
Fragmented
Missing
Complete
Partial
BUSCO
CEGMA

**Figure S4. Visualisation of genome completeness metrics using BUSCO and CEGMA.**
**A.** Comparison of BUSCO completeness among Clade V nematodes. Heatmap presents the complete, fragmented, missing and duplicated BUSCOs of the 982 total BUSCOs (rows) analysed in each genome assembly (columns) of the Clade V nematodes in WBP11. The *H. contortus* V4 genome is indicated in bold. Although BUSCOs are used as a measure of genome completeness, some evidence of phylogenetic structure is present in the column dendrogram, which represents single-linkage clustering of the BUSCO profiles of each species, suggesting systematic missingness due to gene gain and/or loss over evolutionary history. There is also a clear distinction that parasitic species (dark grey circles) have generally poorer BUSCO scores than free-living species (light grey circles). Note this is somewhat confounded by the fact that most free-living species are *Caenorhabditis spp.*, of which *C. elegans* was one species used to develop the BUSCO gene set. **B.** Comparison of genome completeness metrics for all Wormbase Parasite Release 11 species. BUSCO and CEGMA data were obtained for each species from the Wormbase Parasite [5] API, including the *H. contortus* V4 genome (red point).

#### Section 2. Resolving haplotypic diversity and repeat distribution within the chromosomes

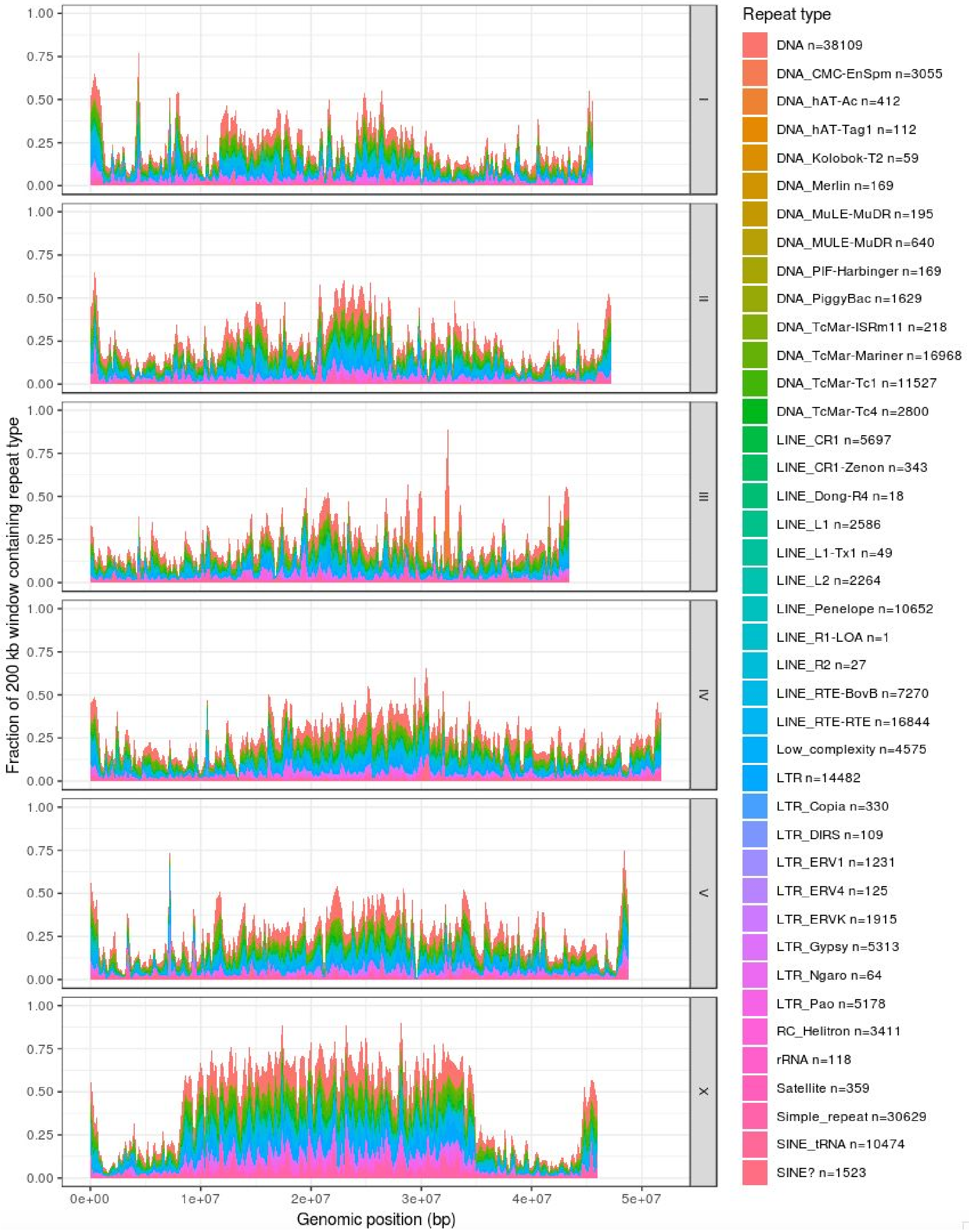


**Figure S5. Distribution of repetitive units throughout the genome identified with RepeatMasker.**

#### Section 3: Generation of a high-quality transcriptome annotation incorporating short and long reads

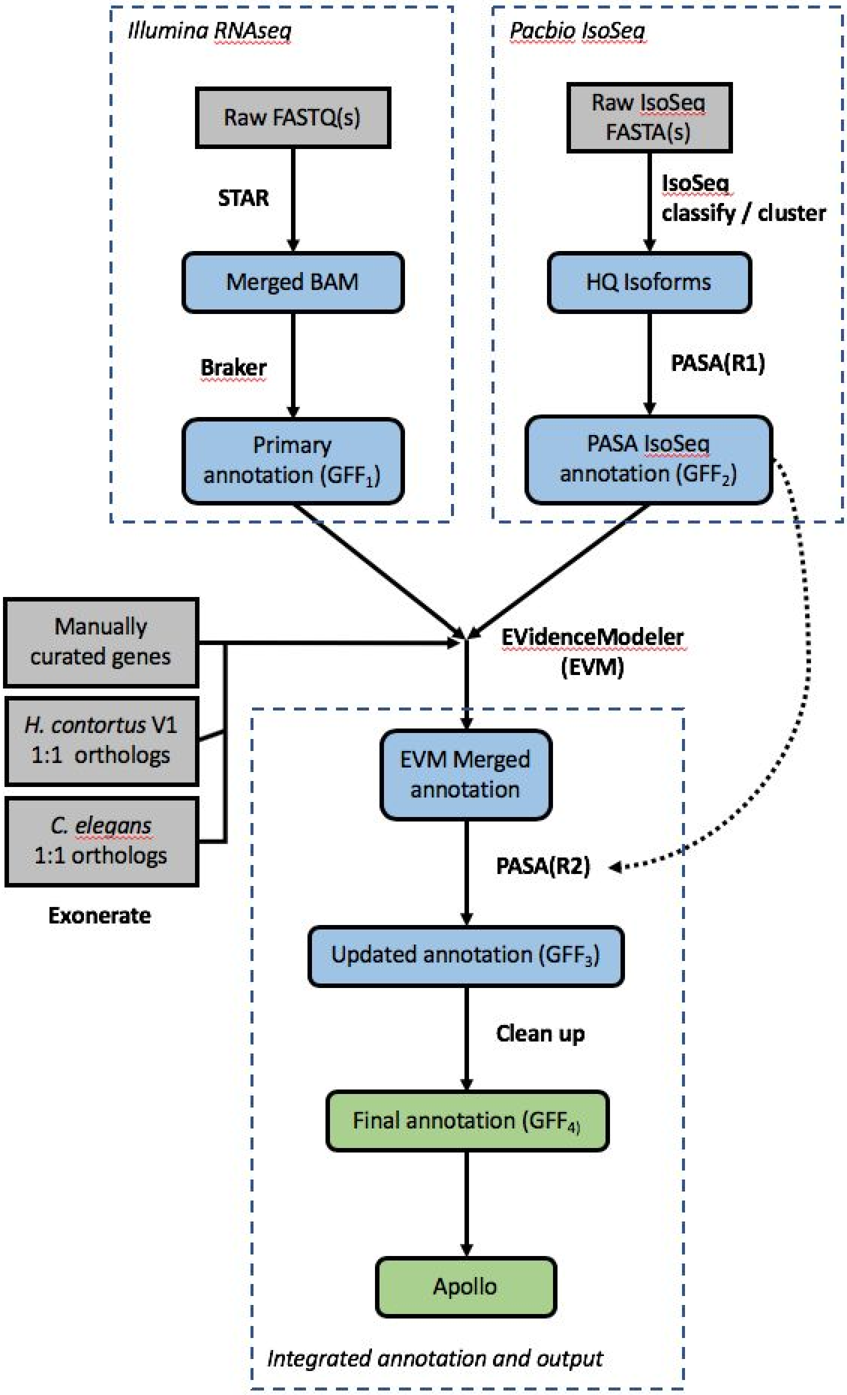


**Figure S6. Annotation pipeline schematic used to incorporate RNAseq and IsoSeq data into a single annotation.**

#### Section 4: Transcriptional dynamics throughout development and between sexes

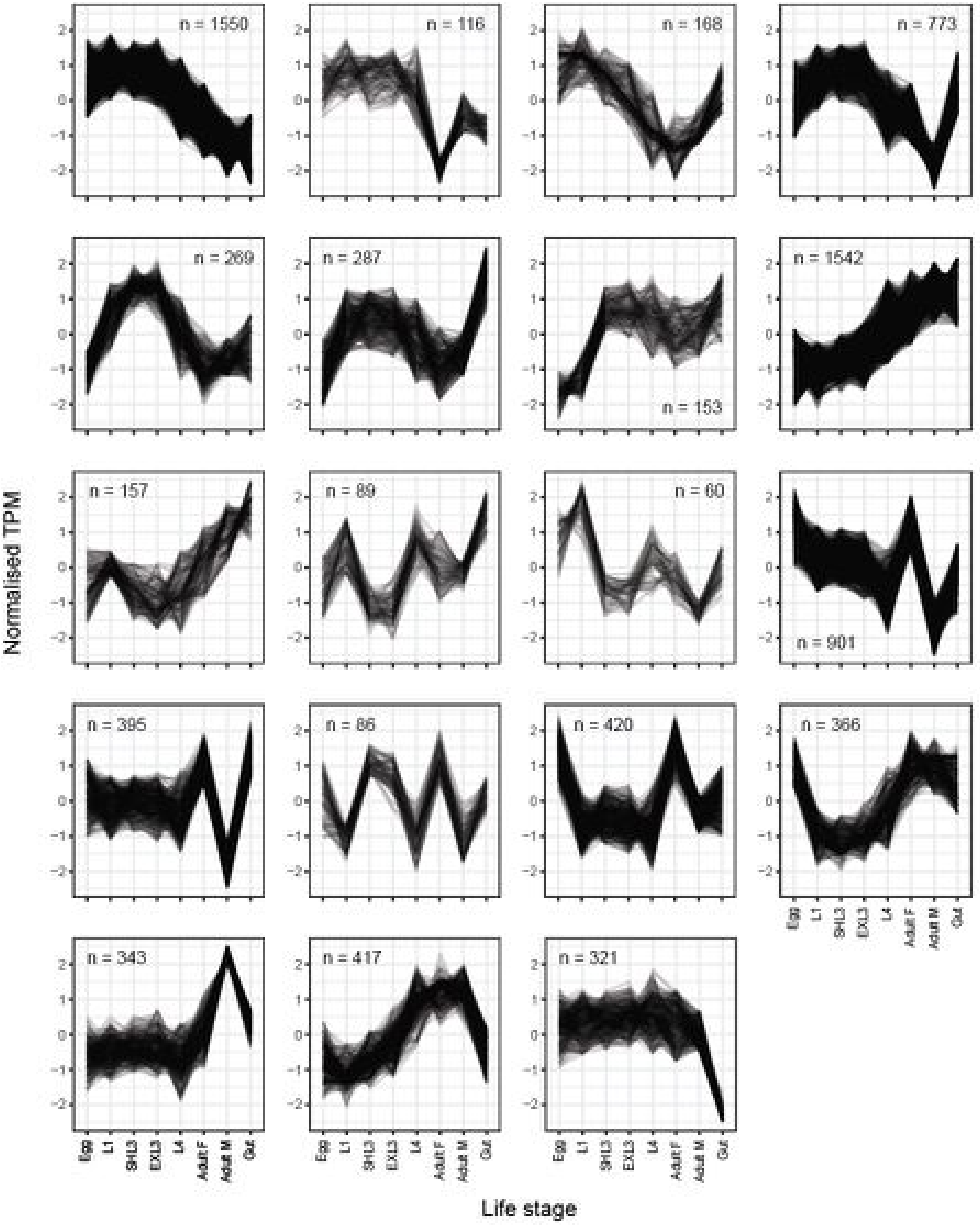


**Figure S7. Transcript co-expression profiles across life stages of *Haemonchus contortus*.** Transcript abundance, defined here as Transcripts per Million (TPM), was determined using Kallisto [6]. Automated clustering of 20986 transcripts using *clust* [7] identified 19 clusters comprising a total of 8412 transcripts with significant but shared differential expression across the life stages, indicative of putative co-regulated transcripts.

#### Section 5: Transcriptional complexity is defined by extensive cis- and trans-splicing

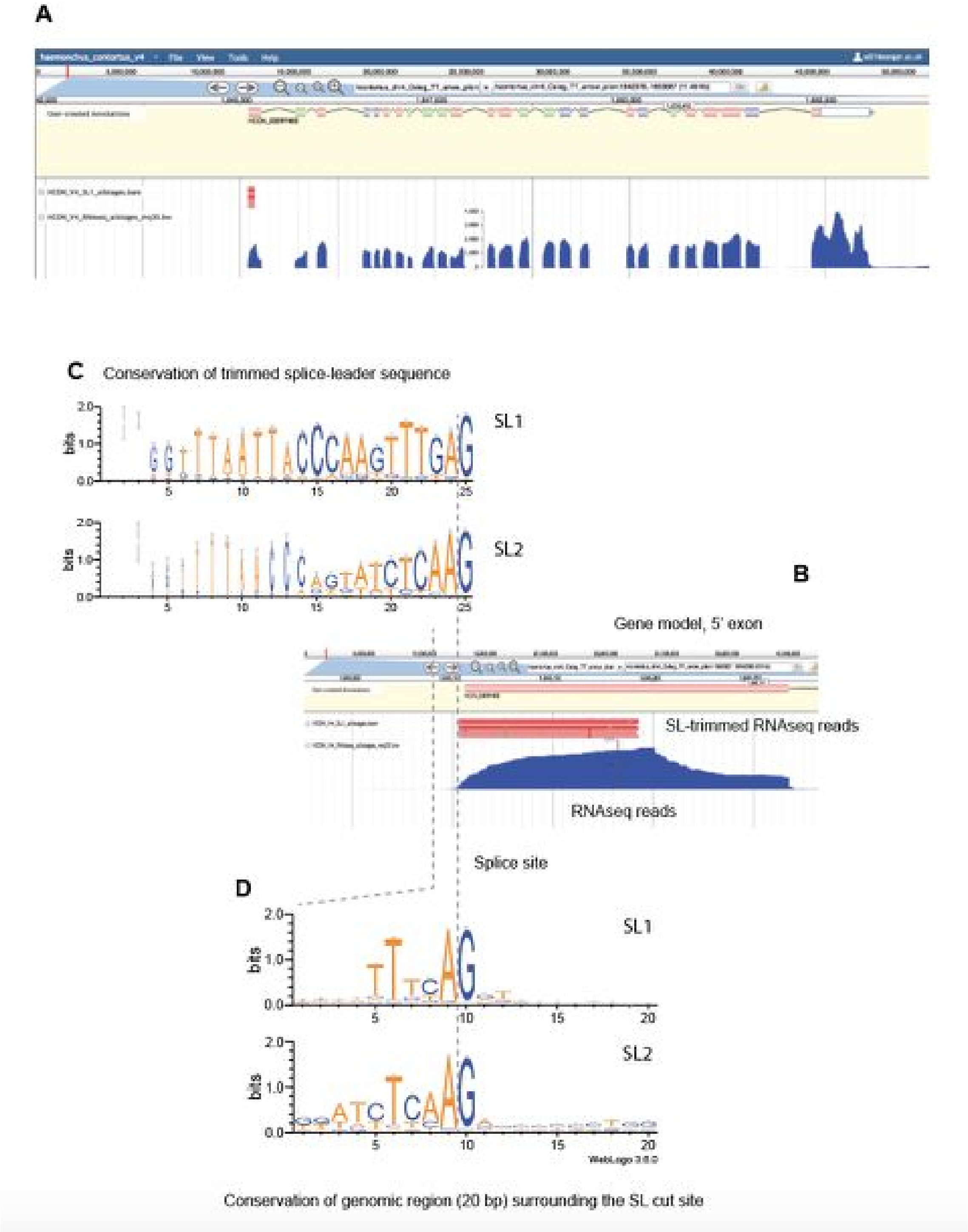


**Figure S8. Comparison of splice leader and splice site sequence diversity**

**A.** WebApollo screenshot of the HCON_00097400 gene model, SL1-trimmed RNAseq reads (red), and RNAseq data supporting the gene model. **B.** Zoomed view of the 5' end of the gene model, highlighting the hard-trimmed RNAseq reads indicative of a match and removal of the SL1 sequence at the same genomic coordinates. This represents the splice site. **C.** Conservation of the DNA sequence containing the trimmed SL1 or SL2 sequence, visualised using WebLogo. The decrease in sequence conservation upstream and away from the splice site is largely due to the decrease in RNAseq read coverage as a result of mRNA decay. **D.** Conservation of genomic DNA sequence surrounding the splice site coordinates, demonstrating the high conservation of AG splice site, but also, enrichment in sequence motifs that differ between SL1- and SL2-targeted genes.

#### Section 5: Distribution of global genetic diversity throughout the chromosomes

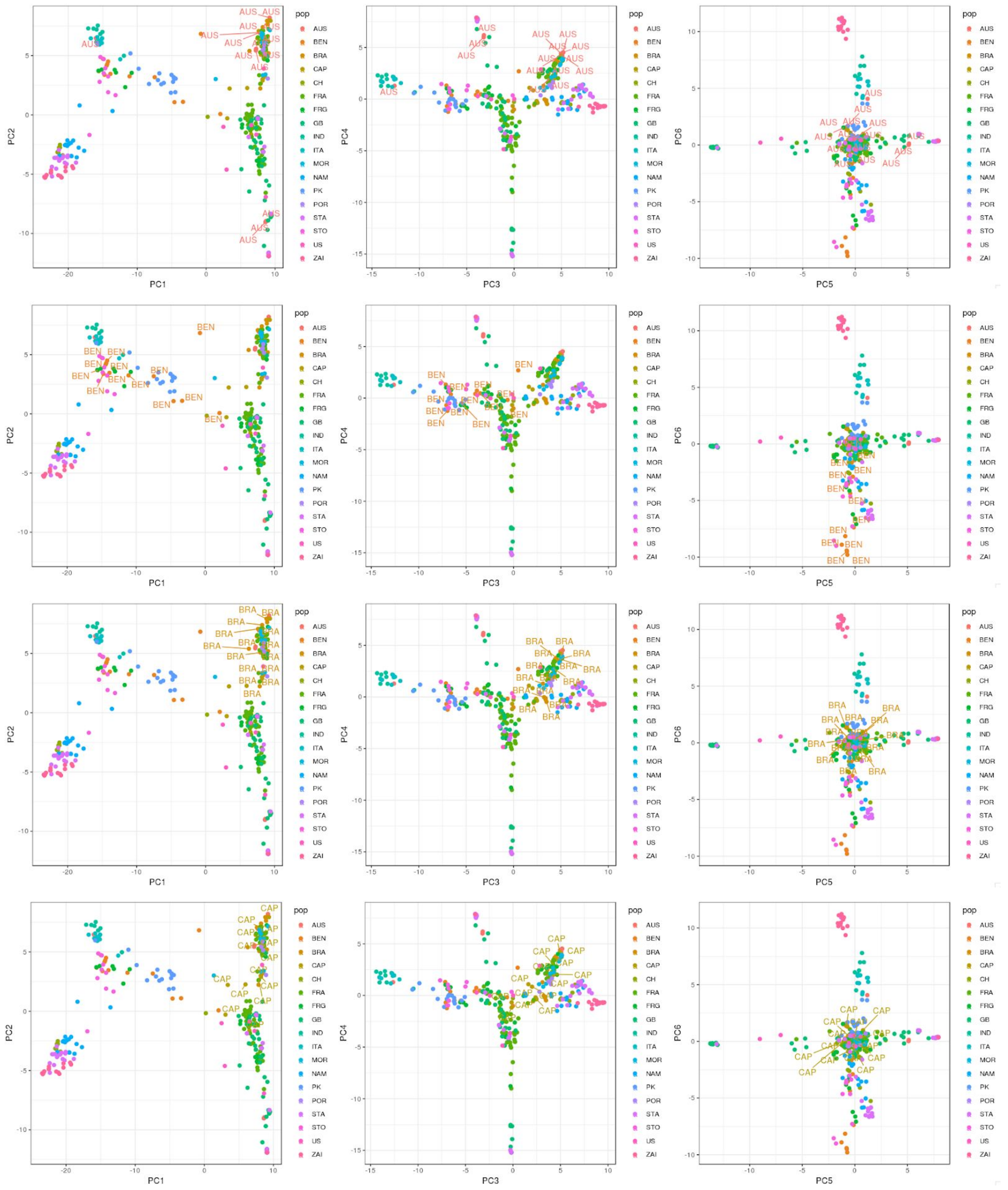

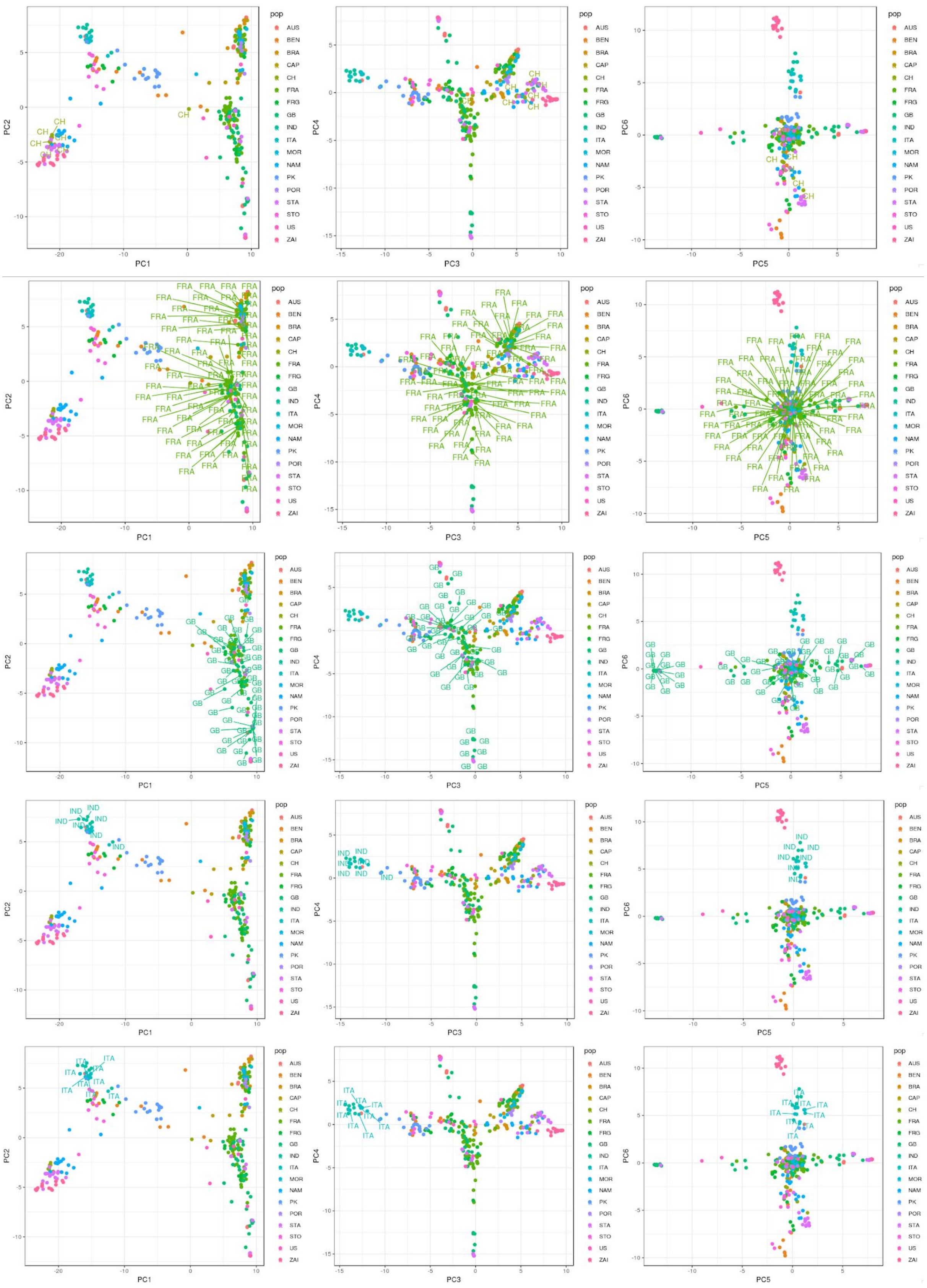
PC1
PC2
PC3
PC4
PC5
PC6
pop
AUS
BEN
BRA
CAP
CH
FRA
FRG
GB
IND
ITA
MOR
NAM
PK
POR
STA
STO
US
ZAI

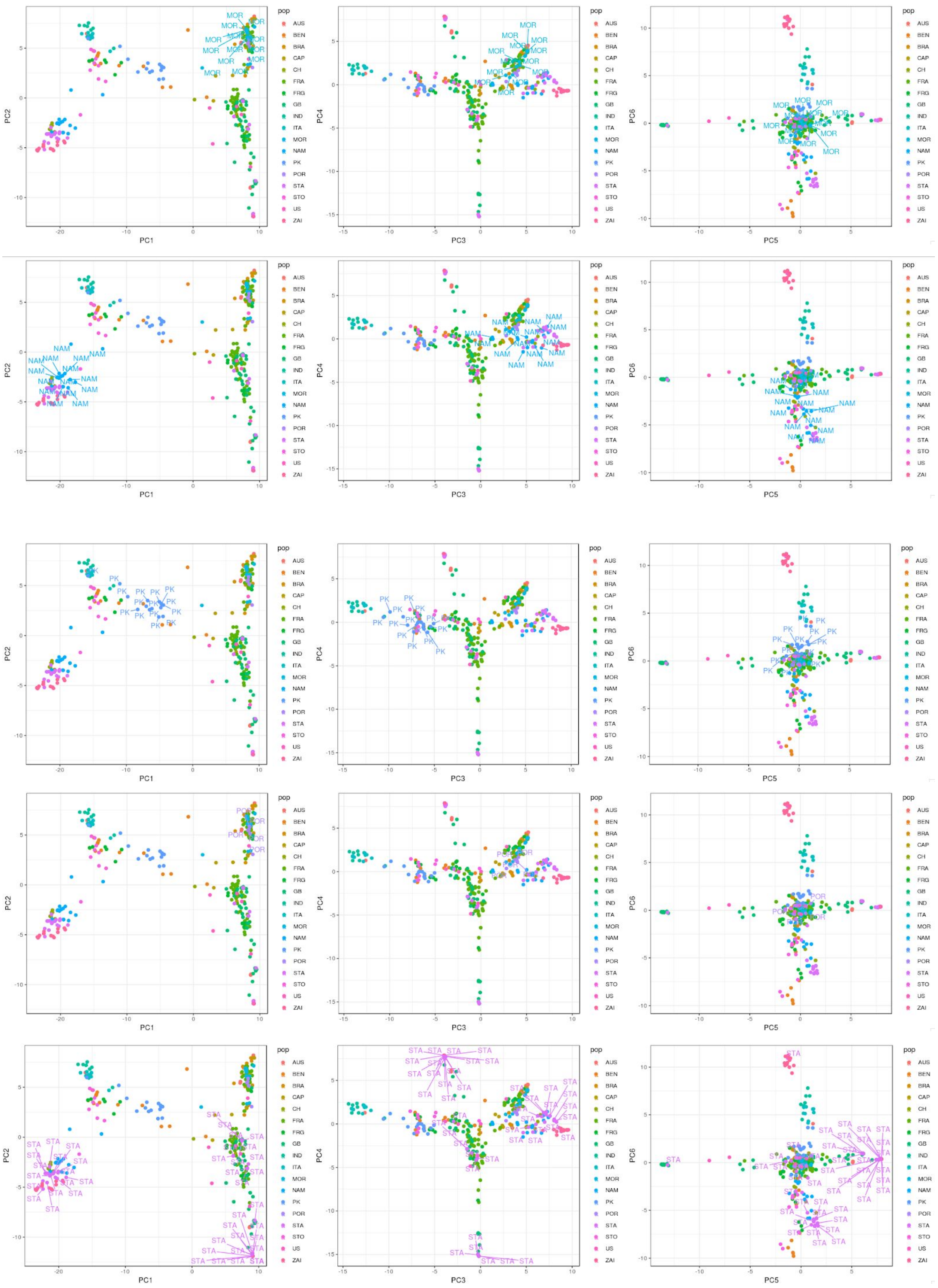
PC1
PC2
PC3
PC4
PC5
PC6
pop
AUS
BEN
BRA
CAP
CH
FRA
FRG
GB
IND
ITA
MOR
NAM
PK
POR
STA
STO
US
ZAI

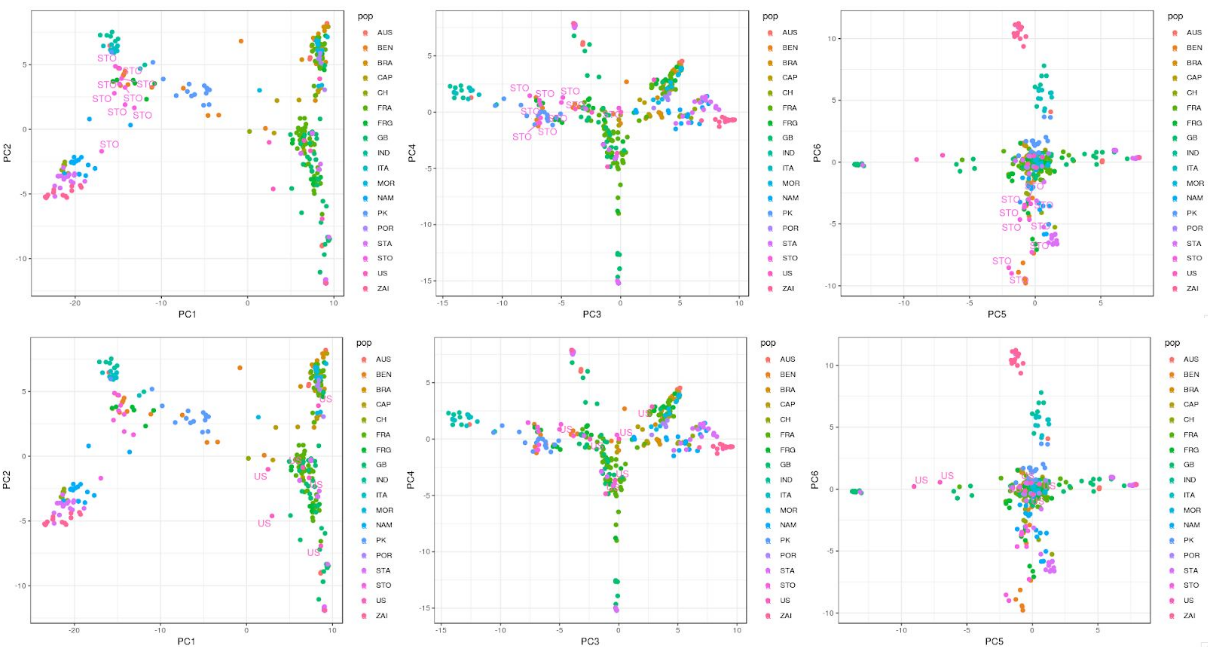


**Figure S9. Principal component analysis (PCA) of mitochondrial diversity**
We extend the analysis of genetic diversity shown in Figure 6 B of the main text to show additional principal component comparisons (PC1v2, PC3v4, PC5v6) and labelling to highlight clustering of samples by country. Countries: Australia (AUS), Benin (BEN), Brazil (BRA), Cape Verde (CAP), Switzerland (CH), France (FRA), Guadeloupe (FRG), United Kingdom (GB), Indonesia (IND), Italy (ITA), Morocco (MOR), Namibia (NAM), Pakistan (PK), Portugal (POR), South Africa (STA), Sao Tome (STO), and United States (US).

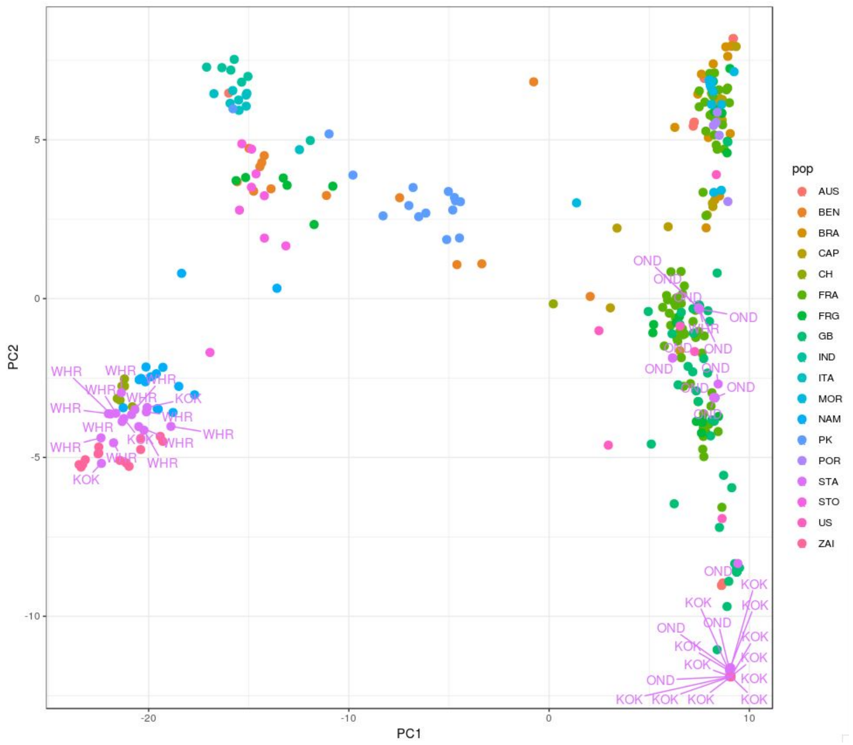


**Figure S10. Distinct differences between mtDNA diversity in South Africa**
Three South African populations of parasites are included, White River (WHR), Kokstadt (KOK), and Onderstepoort (OND). Despite their close proximity geographically, they are genetically distinct.

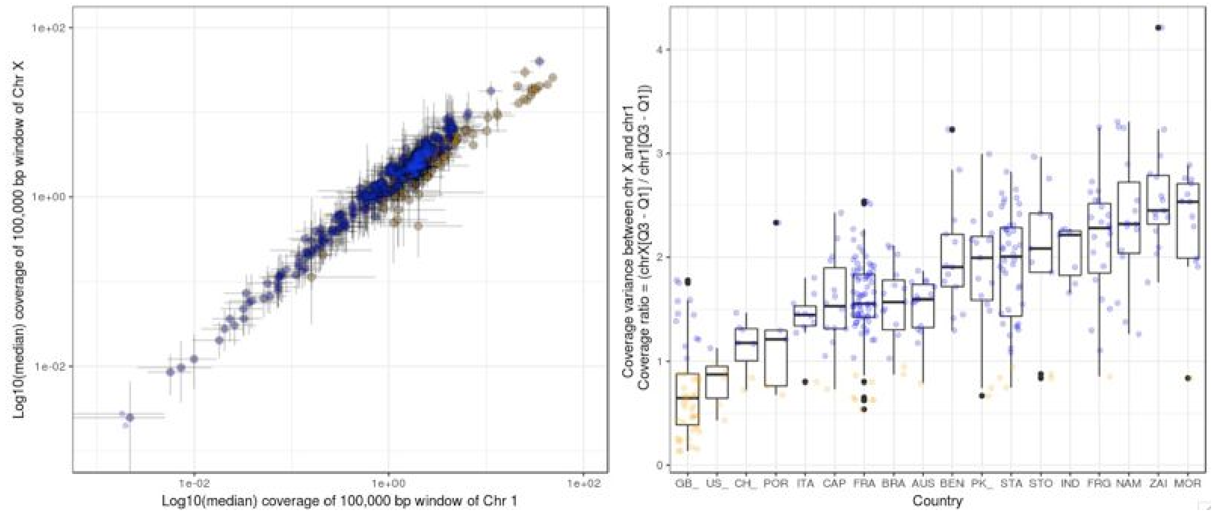


**Figure S11. X chromosome coverage variation**
**A.** Coverage variation between the X chromosome and chromosome I per sample. Error bars represent the 1st (min) and 3rd (max) quartiles of the coverage distribution for the X (y-axis) and autosome (x-axis). **B.** Coverage variation of the X chromosome, normalised by the coverage on the autosomes, per population. Populations are ordered by median coverage from lowest median variation to highest.
